# Pleasure in groove is associated with neuromelanin levels in the substantia nigra of younger healthy individuals

**DOI:** 10.1101/2025.05.10.653047

**Authors:** Takahide Etani, Shinichiro Nakajima, Shiori Honda, Saki Homma, Yuka Kaneko, Sotaro Kondoh, Ryosuke Tarumi, Sakiko Tsugawa, Sotaro Moriyama, Yui Tobari, Tomohiro Samma, Guillermo Horga, Clifford Cassidy, Hiroyuki Uchida, Shinya Fujii

**Affiliations:** Graduate School of Media and Governance, Keio University, Kanagawa, Japan; Keio Research Institute at SFC, Kanagawa, Japan; Keio University Hospital, Tokyo, Japan; Japanese Red Cross Ashikaga Hospital, Tochigi, Japan; Department of Neuropsychiatry, Keio University School of Medicine, Japan; Department of Psychiatry, University of Toronto, Toronto, ON, Canada; Department of Psychiatry and Behavioral Health, Renaissance School of Medicine at Stony Brook University, Stony Brook, NY, USA; Faculty of Policy Management, Keio University, Fujisawa, Kanagawa, Japan; Faculty of Environment and Information Studies, Keio University, Kanagawa, Japan; Japan Society for the Promotion of Science, Tokyo, Japan; Seikei-Kai Komagino Hospital, Tokyo, Japan; Department of Psychiatry, Columbia University

**Keywords:** groove, move, pleasure, music, dopamine, neuromelanin, substantia nigra

## Abstract

The pleasurable urge to move in response to music is called groove. Prior research has suggested a potential link between groove and dopamine function. However, no studies to date have directly investigated the relationship between the two. Here, we aimed to assess individual dopamine function in the substantia nigra of healthy individuals using neuromelanin-sensitive magnetic resonance imaging (NM-MRI), a non-invasive method associated with dopamine function, and to investigate the relationship between the individual dopamine proxy index and sensitivity to the groove experience. In this study, 15 younger (< 48 years) and 16 older (≧48 years) healthy individuals participated. Participants listened to ten musical excerpts and rated the groove experience based on “pleasure” and “wanting to move.” To assess whether the groove experience is related to NM levels, type of musical excerpts, and sex, we analyzed with linear mixed-effects regression models. The results showed that higher NM levels (*p* = 0.032) and male sex (*p* = 0.034) were associated with higher pleasure ratings in the younger group. For the “urge to move” ratings, type of musical excerpts was associated with ratings in both groups (*ps* < 0.001), where high-groove music (Janata et al., 2012) receiving higher ratings. Taken together, these results suggest that the “pleasure” aspect of the groove experience in younger individuals was related to dopamine levels in the substantia nigra, but may not be associated with the “urge to move.” Thus, pleasure and the urge to move are likely to involve distinct dopaminergic pathways and mechanisms, warranting further investigation.

## 1. Introduction

When we listen to music, there are moments when we feel the urge to move our body to the rhythm, such as by bobbing our head, clapping our hands, or stomping our feet. This “pleasurable sensation of wanting to move to music” is called groove (Etani et al. 2024; Stupacher et al. 2016; Witek et al. 2014), or more recently, PLUMM (the abbreviation of the “PLeasurable Urge to Move to Music”) (Matthews et al. 2023), which distinguishes the sensation from a broader concept of groove (Duman et al. 2023). Early experimental research, mainly in the field of psychology, on groove has demonstrated the musical features and individual traits associated with groove, including elements such as syncopation (Witek et al., 2014; Zalta et al., 2024), event density (Madison et al. 2011), tempo (Etani et al. 2018; Jerjen et al. 2024), harmony (Matthews et al. 2019), and bass (Stupacher et al. 2016), as well as factors such as age (Cameron et al. 2022) and musical preference (Senn et al. 2018).

A recent study showed that individuals with specific musical anhedonia, who have a selective reduction of musical pleasure without the impairment of musical ability or general anhedonia, experience less pleasure when listening to groove rhythms compared to those without musical anhedonia (Benson et al. 2024). Additionally, patients with Parkinson’s disease, who have dysfunction in dopamine system which leads to difficulty in initiating movement, experience groove differently compared to healthy individuals. While healthy individuals experience the strongest groove when listening to rhythms with moderate levels of syncopation, and less groove when listening to low or high levels of syncopation (i.e., an inverted U-shaped relationship between groove and syncopation), patients with Parkinson’s disease taking dopamine agonists experience a similar degree of pleasure and urge to move across rhythms with varying levels of syncopation (i.e., a flattened pattern) (Pando-Naude et al. 2024). These findings suggest that dopamine function may be related to individual differences in experiencing groove.

Advancements in neuroscientific research on groove have recently revealed its neural correlates. For instance, using transcranial magnetic stimulation (TMS), one study noted that high-groove music increased corticospinal excitability in musicians compared to low-groove music (Stupacher et al. 2013), suggesting that high-groove music engages cortico-spinal pathways even without physical movement. Furthermore, research using functional magnetic resonance imaging (fMRI) found that listening to high-groove music activates reward and motor-related areas such as the nucleus accumbens (NAcc) and supplementary motor area (Matthews et al. 2020). This finding indicates that the groove experience may be related to dopamine function, which plays a critical role in experiencing pleasure as well as in initiating and controlling movement. Notably, other evidence supports the role of dopamine function in experiencing pleasure while listening to music. A study by Salimpoor et al. (2011) revealed that anticipation and the experience of pleasure while listening to music are associated with dopamine release in the caudate and NAcc, respectively. Furthermore, Ferreri et al. (2019) demonstrated that listening to music enhanced the pleasure experience following the administration of a dopamine agonist (L-Dopa), whereas pleasure decreased after taking a dopamine antagonist (risperidone). These findings further emphasize the role of dopamine in experiencing pleasure from music and support its functional involvement in the groove experience, particularly in relation to the pleasurable aspects of groove. However, while these studies point to the involvement of the dopamine system in the groove experience, no previous research has directly investigated this relationship.

For many years, positron emission tomography (PET) has been used as a powerful tool to quantify dopamine function in vivo using radioactive tracers. However, dopamine PET is not widely applied outside the research settings because of its invasive and costly nature. Recently, neuromelanin (NM)-sensitive MRI (NM-MRI) has been developed to quantify dopamine function indirectly and noninvasively (Cassidy et al. 2019). NM is a pigment that accumulates in neurons, particularly dopaminergic neurons in the substantia nigra (SN) (Zucca et al. 2014), and has been shown to correlate with PET measures that reflect dopamine function (Cassidy et al. 2019). It has been reported that NM levels change with age in healthy individuals and display different patterns in patients with Parkinson’s disease and schizophrenia, both of which involve dysfunctions in the dopamine system (Ueno et al. 2022). This suggests that measuring NM levels is useful for quantifying individual differences in dopamine function. Furthermore, while PET assesses short-term dopamine function, NM levels reflect long-term dopamine function. Therefore, NM-MRI signal may more closely reflect trait-like, rather than state-like, characteristics in each individual.

Therefore, the aim of this study was to examine the relationship between dopamine function and individual sensitivity to the groove experience. We also sought to investigate whether the groove experience may differ depending on musical excerpts as well as sex. For these aims, we conducted a cross-sectional study measuring the NM levels in the SN using NM-MRI and collecting groove ratings for musical excerpts during a listening experiment in healthy individuals. We hypothesized that groove ratings would be related to NM levels in the SN of healthy individuals, where healthy individuals who have higher NM levels (indicating higher dopamine function) would experience stronger groove when listening to music. Additionally, we hypothesized that healthy individuals would experience stronger groove with musical excerpts that had received high groove ratings in a previous study (Janata et al. 2012), thereby validating the selection of stimuli used in this study. Furthermore, we hypothesized that female participants would experience stronger groove since females are reported to have higher sensitivity to musical reward (Mas-Herrero et al. 2012; Honda et al. 2023).

## 2. Method

### 2.1 Overview

This single-center cross-sectional study was conducted at Komagino Hospital, Tokyo, Japan, in compliance with the Declaration of Helsinki. Data analysis was conducted at Keio University School of Medicine, Tokyo, Japan and Komagino Hospital, with approval granted by the ethics committees of these constitutions (approval numbers: 20170313 and 20230003).

### 2.2 Participants

The inclusion criteria for healthy individuals are no history of psychiatric illness confirmed by qualified psychiatrists. The exclusion criteria were as follows: 1) active substance abuse or dependence during the past 6 months, 2) positive urine drug screen for drugs of abuse at inclusion or before MRI scanning, 3) history of head trauma resulting in loss of consciousness for >30 minutes, 4) unstable physical illness, or 5) neurological disorder. All participants provided written informed consent for this study before enrollment. They were recruited from the local community in Tokyo, Japan. The sample size was calculated for a parental study to investigate the relationship between SN NM levels and treatment responsiveness in schizophrenia. Based on an effect size reported from a previous study that compared SN NM levels in patients with schizophrenia to healthy controls (Cohen’s d = 0.92) (Cassidy et al., 2019), which resulted in the estimated number of 24 individuals in each group (alpha = 0.05, 1-beta = 0.85). Since we continued to enroll 24 pairs of each group, we included 31 healthy individuals (mean age = 45.8 ± 12.4 years, 12 females) participated in the study. We divided the participants into two groups: a younger group and an older group (Table 1). Details are provided in Section 2.8.

**Table 1.**
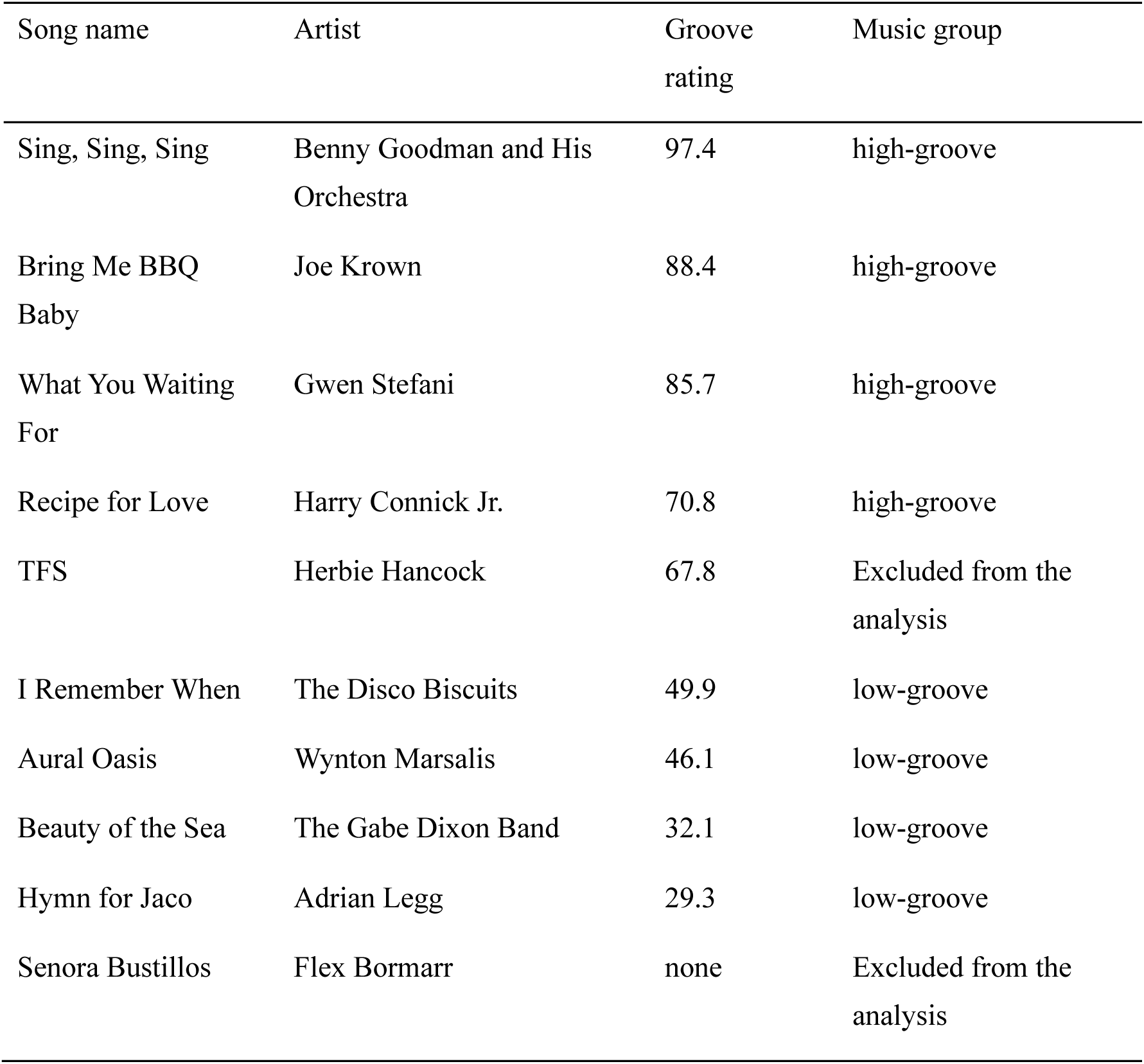
Musical stimuli used in the study. Nine of them were selected from a previous study (Janata et al., 2012). One was selected as a control stimulus.

### 2.3 Stimuli

In the study, we used ten musical excerpts as stimuli. Nine of these were selected from a previous study that obtained groove ratings (Janata et al. 2012), and one (‘Senora Bustillos’ by Flex Bormarr) was selected as a control stimulus that was not included in Janata et al (2012). We extracted 20-second segments from each (Table 2). Musical excerpts selected from Janata et al. (2012) included a variety of music styles, with groove ratings ranging from 0 to 128 in the original study. To ensure a broad range of groove levels, we selected excerpts with a wide range of ratings, specifically those ranging from 29.3 to 97.4. Out of the ten songs used in the experiment, we excluded one musical excerpt from the analysis because it was not used in the previous study (Janata et al. 2012) and was included in our study as a control stimulus. We also excluded one musical excerpt with a mid-groove rating in the previous study (Janata et al. 2012) in order to include an equal number of high-groove and low-groove musical excerpts in the analysis. Eventually, we included eight (i.e., four high-groove and four low-groove) musical excerpts in the analysis, and grouped them into two categories; one with high groove ratings (high-groove excerpts) and the other with low groove ratings (low-groove excerpts), based on the results of (Janata et al. 2012).

**Table 2.** Demographic data of the participants.

| | Younger group ($n = 14$) | Older group ($n = 16$) |
| --- | --- | --- |
| Age, years | $34.36 \pm 8.14$ | $55.81 \pm 4.99$ |
| Sex, female | 7 (50.0%) | 5 (31.3%) |

### 2.4 Procedure

Participants listened to ten musical excerpts (each 20 seconds long) through headphones (Bose, USA) and rated “How much pleasure did you feel?” (hereinafter referred to as “pleasure” ratings) and “How much did you feel like moving your body?” (hereinafter referred to as “wanting to move” ratings) on a five-point Likert scale using an application (Qualtrics, USA) on an iPad. Following this, we measured each participant’s NM levels using a 3T MRI scanner (GE, USA). The region of interest was the SN in the midbrain.

### 2.5 Magnetic resonance imaging

MRI data were acquired for all participants using a 3T GE Signa HDxt scanner with an 8-channel head coil. Participants underwent a 3D inversion recovery-prepared T1-weighted MRI scan (Axial MRI 3D brain volume [BRAVO]; repetition time [TR] = 6.4 ms, echo time [TE] = 2.8 ms, inversion time [TI] = 650 ms, flip angle = 8°, field of view [FOV] = 230 mm, matrix = 256 × 256, and slice thickness = 0.9 mm).

### 2.6 NM-MRI acquisition and data processing

NM-MRI images were acquired using a two-dimensional gradient echo sequence with magnetization transfer contrast (TR = 260 ms, TE = 3.72 ms; flip angle = 40°, in-plane resolution = 0.43 x 0.43 mm2, partial brain coverage with FOV = 220 x103, matrix 240 x 512, number of slices = 8, slice thickness = 3 mm, slice gap = 3 mm, magnetization transfer frequency offset = 1200 Hz, number of excitations = 8, and acquisition time = 17.88 min). The protocol for slice prescription involved aligning the image stack along the line between the anterior and posterior commissures, with the top slice positioned 3 mm above the floor of the third ventricle. This protocol covered the SN-containing portions of the midbrain and surrounding structures. The quality of the NM-MRI images was inspected visually.

During preprocessing, we used ANTs (Avants et al., 2008) and SPM12 running on MATLAB (R2022b). Preprocessing steps were carried out in accordance with the methodologies described in previous studies (Wengler et al., 2020; Cassidy et al., 2019). NM-MRI scans were co-registered to each participant’s T1-weighted anatomical image and normalized to MNI space using ANTs. All images were visually inspected after each preprocessing step to ensure quality. Subsequently, intensity normalization and spatial smoothing were applied sequentially using custom MATLAB scripts. The contrast-to-noise ratio (CNR) for each participant and voxel(_*v*_) was calculated as the relative difference in NM-MRI signal intensity (*I*) from a reference region (*CC*) in the white matter—specifically, the crus cerebri, which is known to contain minimal neuromelanin— according to the following formula:

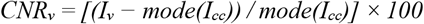

The reference region mask for the SN was derived from a previous study (Cassidy et al., 2019), and the SN mask itself was adapted from the same source with minor modifications. These masks, overlaid on the group-averaged CNR image, are presented in Figure 1. For each participant, the mode (referred to as *I*_*CC*_) was computed by applying a kernel-smoothing function to the histogram of voxel intensity values within the mask. The resulting NM-MRI CNR maps were then further smoothed using a Gaussian kernel with a full width at half maximum (FWHM) of 1 mm. Finally, the average of CNR_*v*_ was calculated for each participant from all voxels in the SN mask. A map of the SN and reference region masks used in the calculation of CNR is shown in Figure 1. Data from one participant were excluded from the statistical analysis due to improper mask placement.

**Figure 1.**
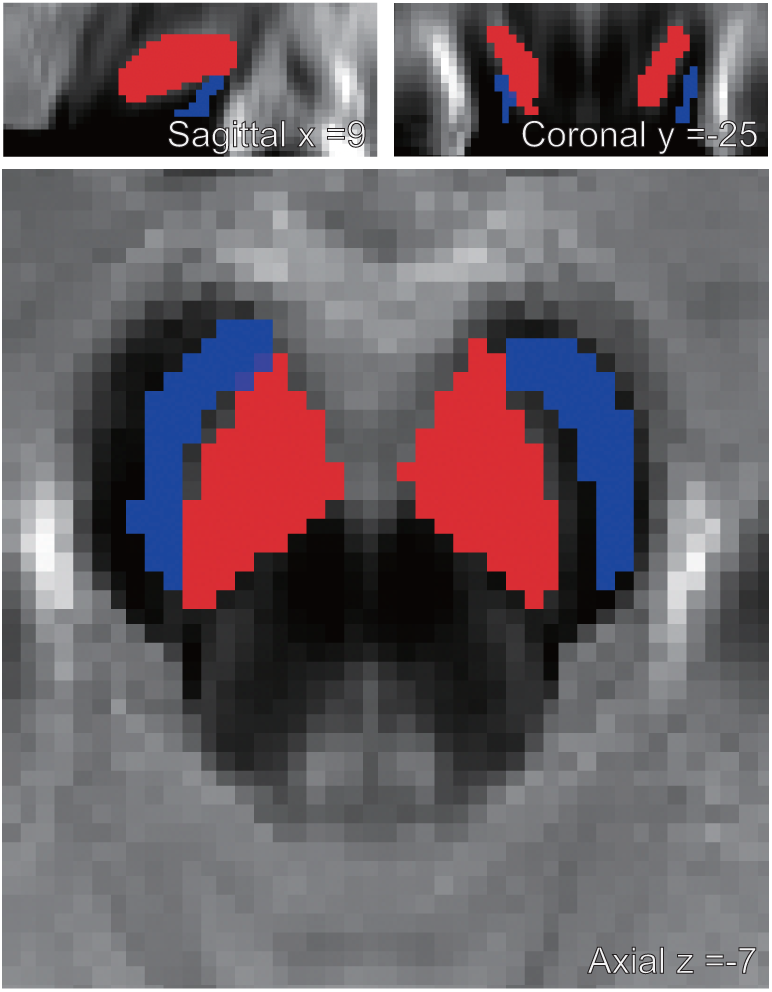
A map of voxels in the substantia nigra (SN) and crus cerebri (CC) used in the calculation of Neuromelanin-MRI contrast-to-noise ratio (CNR). The masks for the SN (red voxels) and the CC (blue voxels) are overlaid onto the template.

### 2.7 Statistical analysis

Statistical analyses were performed using R version 3.6.3. First, we calculated the average “pleasure” ratings as well as the “wanting to move” ratings for the four musical excerpts in high-groove excerpts for each participant. Similarly, we calculated the average “pleasure” ratings and the average “wanting to move” ratings for the four musical excerpts in low-groove excerpts for each participant.

Next, we divided participants into two groups based on a previous study (Xing et al. 2018): a younger group (under 48 years old, *n* = 14) and an older group (48 years and older, *n* = 16), as the profile of NM level varies with age (Xing et al. 2018). They found an inverted U-shaped relationship between NM levels and age, where NM levels rapidly accumulate up to 20 years of age, followed by a slow increase and a plateau between approximately 45 and 52 years, and then a decrease with further aging. Following this pattern, they divided the population into three groups: under 20 years, 21 to 47 years, and over 47 years. We followed this study and divided the participants into two groups. Since all the participants were over 20 years old, we did not include the third group (under 20 years old). Next, we performed the Kolmogorov-Smirnov test for all data (“pleasure” and “wanting to move” ratings for both the younger and older groups) and confirmed that they were all normally distributed.

In order to test whether NM level and music group (high-groove excerpts and low-groove excerpts) are associated with groove ratings, we constructed two linear mixed-effects regression models with “pleasure” and “wanting to move” ratings as the response variables, NM-MRI CNR, music group (high-groove excerpts and low-groove excerpts), and sex (male and female) as fixed effects, and participant ID as a random effect for the younger group. Similarly, we constructed the same linear mixed-effects regression models for the older group. The significance level was set at *p* < 0.05 for each analysis.

## 3. Results

The demographic data is shown in Table 2.

### 3.1 Younger group

For the younger group, the results of the linear mixed-effects regression model on the “pleasure” ratings revealed that NM-MRI CNR and sex were associated with “pleasure” ratings. Specifically, individuals with higher NM-MRI CNR in the SN experienced greater pleasure from music, and male participants reported stronger pleasure compared to female participants (*R*^2^_m_ = 0.38, *R*^2^_c_ = 0.74, Table 3). Figure 2 illustrates the relationship between the pleasure rating, adjusted for fixed effects, and the NM-MRI CNR in the SN.

**Table 3.** The results of the linear mixed-effects regression model on the “pleasure” ratings for the younger group (age < 48 years).

| Variables | $\beta$ | *SE* | *t* value | *p* value |
| --- | --- | --- | --- | --- |
| Slope | 1.80 | 1.36 | 1.32 | 0.21 |
| NM-MRI CNR (%) | 0.22* | 0.089 | 2.46 | 0.032 |
| Sex (1 = male, 2 = female) | −0.77* | 0.32 | −2.42 | 0.034 |
| Music Group (1 = high-, 2 = low-groove) | −0.036 | 0.17 | −0.21 | 0.83 |
| * *p* < 0.05 ** *p* < 0.01 *** *p* < 0.001 |  |  |  |  |

Note: For the variable "Sex," we used dummy coding, assigning 1 to male and 2 to female. For the variable "Music Group," we also used dummy coding, assigning 1 to high-groove music and 2 to low-groove music.

**Figure 2.**
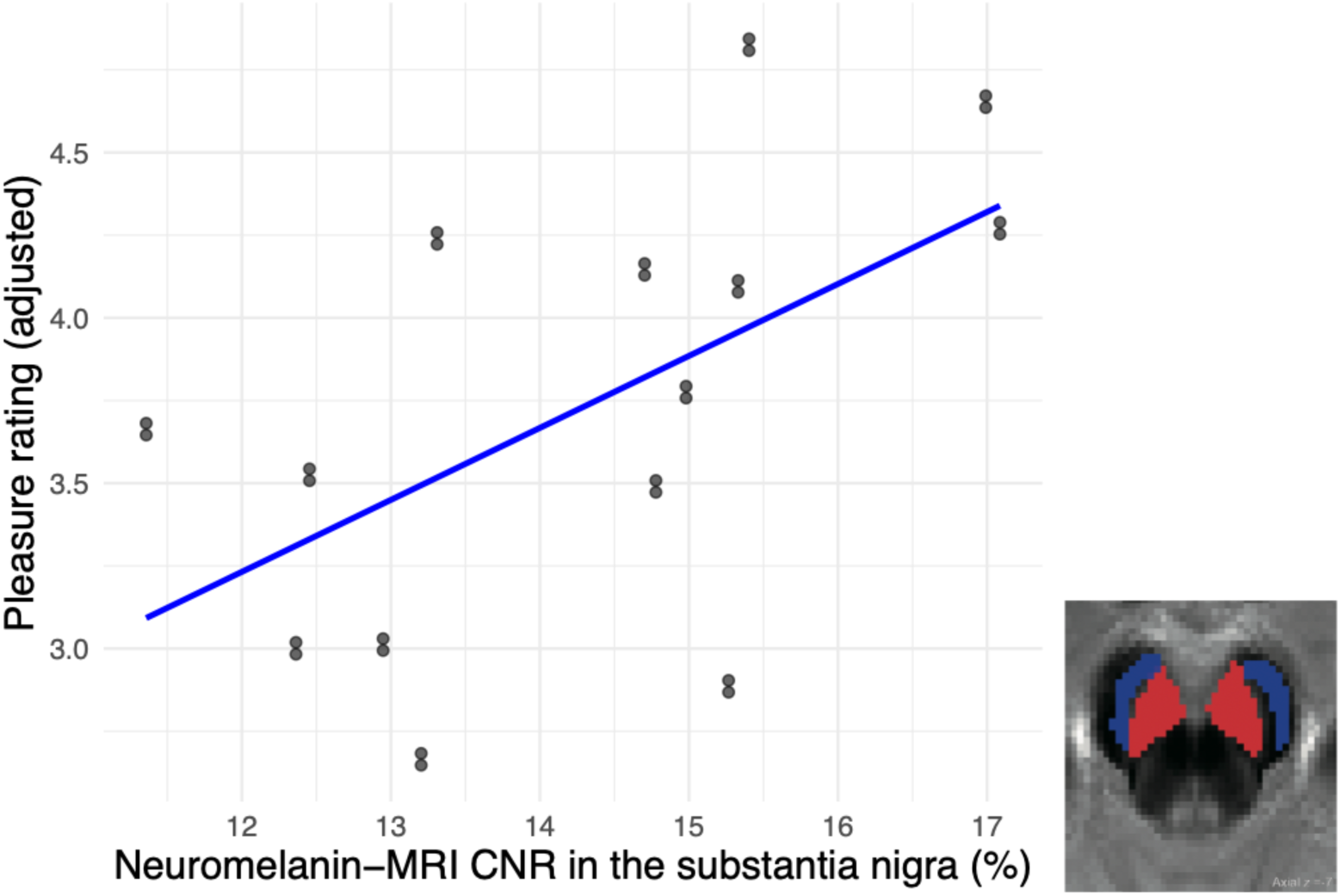
The relationship between the NM-MRI CNR in the substantia nigra and pleasure rating with random effects in the younger group.

The results of the linear mixed-effects regression model on the “wanting to move” ratings revealed that the music group was associated with the ratings. Specifically, high-groove excerpts, which received high groove ratings in the previous study (Janata et al. 2012), elicited higher “wanting to move” ratings compared to low-groove excerpts, which had low groove ratings in the same study (*R*^2^_m_ = 0.45, *R*^2^_c_ = 0.78, Table 4).

**Table 4.** The results of the linear mixed-effects regression model on the “wanting to move” ratings for the younger group (age < 48 years).

| Variables | $\beta$ | *SE* | *t* value | *p* value |
| --- | --- | --- | --- | --- |
| Slope | 2.02 | 2.15 | 0.94 | 0.37 |
| NM-MRI CNR (%) | −0.037 | 0.14 | −0.27 | 0.80 |
| Sex (1 = male, 2 = female) | −0.74 | 0.50 | −1.48 | 0.17 |
| Music Group (1 = high-, 2 = low-groove) | 1.68*** | 0.25 | 6.70 | <0.001 |
| * *p* < 0.05 ** *p* < 0.01 *** *p* < 0.001 |  |  |  |  |

Note: For the variable "Sex," we used dummy coding, assigning 1 to male and 2 to female. For the variable "Music Group," we also used dummy coding, assigning 1 to high-groove music and 2 to low-groove music.

### 3.2 Older group

For the older group, the results of the linear mixed-effects regression model on the “pleasure” ratings revealed that none of the variables were associated with these ratings (*R*^2^_m_ = 0.10, *R*^2^_c_ = 0.80, Table 5).

**Table 5.** The results of the linear mixed-effects regression model on the “pleasure” ratings for the older group (age = 48 years and older).

| Variables | $\beta$ | *SE* | *t* value | *p* value |
| --- | --- | --- | --- | --- |
| Slope | 3.29** | 1.06 | 3.09 | 0.0084 |
| NM-MRI CNR (%) | 0.033 | 0.076 | 0.43 | 0.67 |
| Sex (1 = male, 2 = female) | 0.31 | 0.29 | 1.07 | 0.31 |
| Music Group (1 = high-, 2 = low-groove) | −0.094 | 0.091 | −1.03 | 0.32 |
| *$p < 0.05$ **$p < 0.01$ ***$p < 0.001$ | | | | |

Note: For the variable "Sex," we used dummy coding, assigning 1 to male and 2 to female. For the variable "Music Group," we also used dummy coding, assigning 1 to high-groove music and 2 to low-groove music.

The results of the linear mixed-effects regression model on the “wanting to move” ratings revealed that the music group was associated with the ratings. Specifically, as for the younger group, high-groove excerpts, which received high groove ratings in the previous study (Janata et al. 2012), elicited higher “wanting to move” ratings compared to low-groove excerpts, which had low groove ratings in the same study (*R*^2^_m_ = 0.45, *R*^2^_c_ = 0.57, Table 6).

**Table 6.**
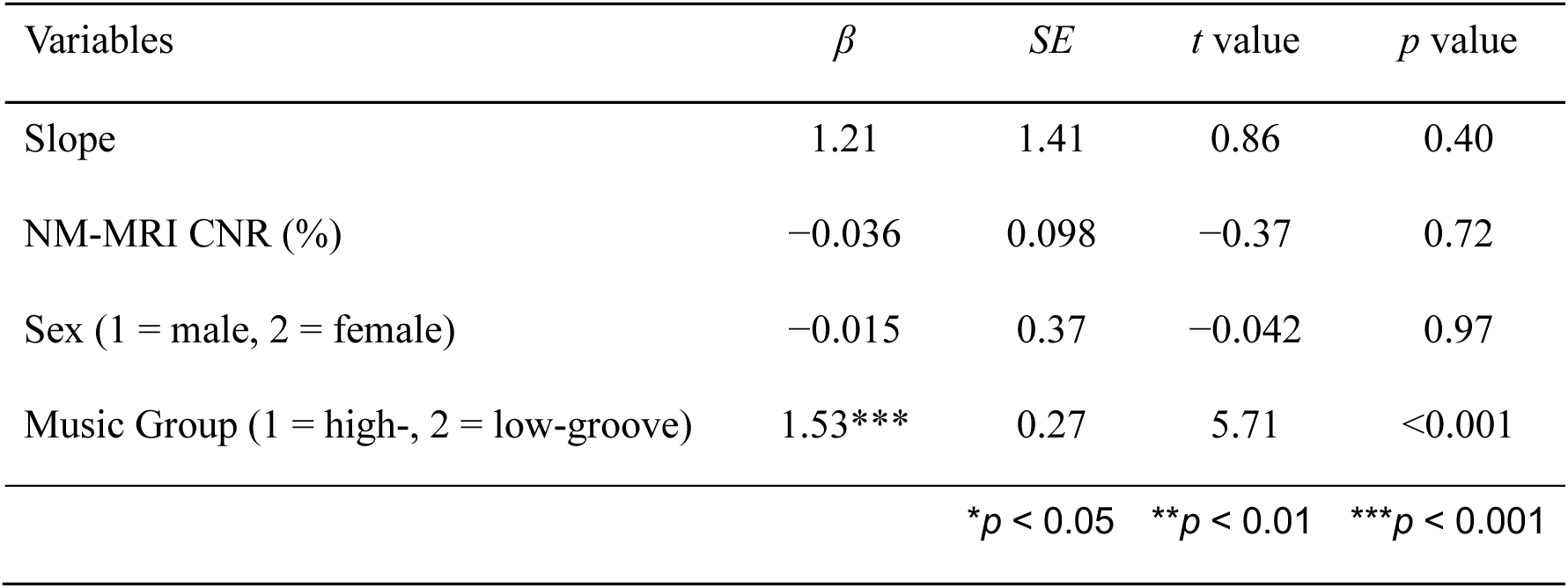
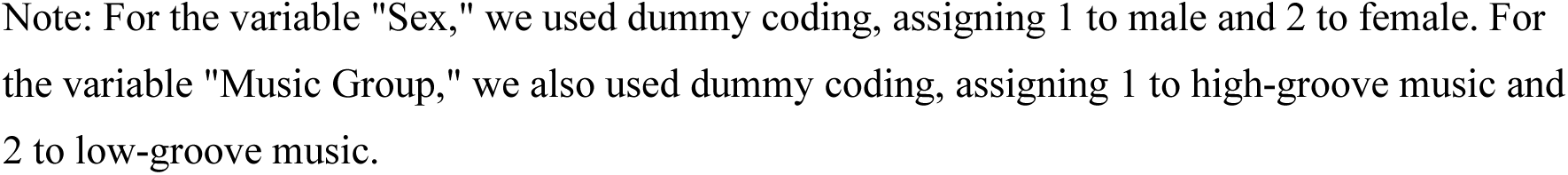
The results of the linear mixed-effects regression model on the “wanting to move” ratings for the older group (age = 48 years and older).

| Variables | $\beta$ | *SE* | *t* value | *p* value |
| --- | --- | --- | --- | --- |
| Slope | 1.21 | 1.41 | 0.86 | 0.40 |
| NM-MRI CNR (%) | −0.036 | 0.098 | −0.37 | 0.72 |
| Sex (1 = male, 2 = female) | −0.015 | 0.37 | −0.042 | 0.97 |
| Music Group (1 = high-, 2 = low-groove) | 1.53*** | 0.27 | 5.71 | <0.001 |
| *$p < 0.05$ **$p < 0.01$ ***$p < 0.001$ | | | | |

Note: For the variable "Sex," we used dummy coding, assigning 1 to male and 2 to female. For the variable "Music Group," we also used dummy coding, assigning 1 to high-groove music and 2 to low-groove music.

## 4. Discussion

In this study, we investigated whether an individual’s NM level measured with MRI (NM-MRI CNR) and type of musical excerpts (high/low-groove musical excerpts), as well as sex affect the groove experience measured with high- and low-groove music in healthy individuals. For the younger group (under 48 years), in terms of the “pleasure” ratings, NM-MRI CNR in the SN and sex were associated with “pleasure” ratings. Specifically, individuals with higher NM-MRI CNR in the SN experienced greater pleasure when listening to music, and male participants reported stronger pleasure compared to female participants. On the other hand, with regard to the “wanting to move” ratings, the music group was associated with the ratings, showing that participants felt a stronger urge to move when listening to high-groove music compared to low-groove music, as identified in the previous study (Janata et al. 2012). For the older group (48 years and older), none of the variables was associated with the “pleasure” ratings. However, for the “wanting to move” ratings, the music group was associated with the ratings, suggesting that, similar to the younger group, participants felt a stronger urge to move when listening to high-groove music compared to low-groove music. The strength of our study lies, first, in being the first to investigate the relationship between dopamine function and the groove experience, revealing an association between dopamine function and the pleasure aspect of groove in young individuals. Second, by applying NM-sensitive imaging, our study was able to assess the relationship between long term, trait-like dopamine function and the groove experience.

We found that NM-MRI CNR in the SN was related to the “pleasure” aspect of groove in the younger group, indicating that individuals with higher dopamine function in the SN experience stronger pleasure when listening to music. It has been shown that dopamine function in the SN activates during periods of heightened reward anticipation (Rios et al. 2023). Therefore, in the younger group, individuals with higher dopamine function in the SN may perceive music as a reward, eliciting strong anticipation and leading to a greater experience of pleasure from music. On the other hand, NM-MRI CNR was not related to the “pleasure” ratings in the older group. Previous research has shown that NM-MRI CNR is positively correlated with PET measures (k_i_^cer^ measured by ^18^F-DOPA PET) in young healthy individuals (mean age = 31.9 years) (Cassidy et al. 2019), indicating that NM-MRI CNR reflects dopamine function in younger populations. However, since NM levels decrease with age (Xing et al. 2018), this decline may diminish the ability of NM-MRI CNR to reflect dopamine function. Therefore, in the older group, NM levels might not have reflected dopamine function as clearly as in the younger group. However, since previous studies have shown that dopamine function decreases with age (Nakajima et al. 2015; Karrer et al. 2017), it is also possible that NM levels also correlate with dopamine function in older populations, suggesting that NM levels in older populations may still reflect dopamine function. In this case, the lack of correlation between pleasure ratings and NM levels could be attributed to other factors, which should be further investigated in future research. In contrast to pleasure, we found no relationship between NM-MRI CNR in the SN and the urge to move. This may be due to different neural pathways being responsible for pleasure and the urge to move, or motivation. While many studies have reported a positive correlation between pleasure and the urge to move (e.g., Düvel et al. 2021; Etani et al. 2018; Senn et al. 2020; Witek et al. 2014), it is important to note that they are distinct aspects of groove. Pleasure is a subjective experience driven by reward processing, whereas the urge to move represents motivation, acting as a trigger to obtain reward or pleasure. Indeed, while pleasure and motivation are both related to dopamine pathways, different pathways are responsible for them (Bressan and Crippa 2005). Pleasure is processed through the mesolimbic pathway, which connects the ventral tegmental area (VTA) in the midbrain to the ventral striatum, including the NAcc in the basal ganglia, as well as throughout the striatum and SN (Haber 2011). Indeed, the experience of pleasure and its anticipation while listening to music are associated with dopamine release in the NAcc and the caudate, respectively (Salimpoor et al. 2011), highlighting the role of the mesolimbic pathway in experiencing musical pleasure. On the other hand, motivation is processed through the mesocortical pathway, which connects the VTA to the prefrontal cortex. Therefore, it is possible that the urge to move is related to dopamine function in the mesocortical pathway, rather than the one associated with the SN.

As we hypothesized, high-groove excerpts, which received high groove ratings in Janata et al (2012), received higher “wanting to move” ratings than low-groove excerpts, which had low groove ratings in the same study. However, contrary to our hypothesis, there was no significant difference in the “pleasure” ratings between high-groove excerpts and low-groove excerpts. These results suggest that participants in the previous study, who were asked to rate “groove” without separating it into “pleasure” and “wanting to move,” associated groove more with the urge to move than with pleasure. This result also highlights the importance of measuring both the urge to move and pleasure to assess the groove experience. It suggests that the two components contribute differently, with the urge to move accounting for a larger proportion of the groove experience than pleasure.

We found that male participants experienced stronger “pleasure” when listening to music. This result is contrary to our hypothesis based on the previous studies using the Barcelona Music Reward Questionnaire (BMRQ) (Mas-Herrero et al. 2012; Honda et al. 2023), which showed that female participants had higher total BMRQ scores (i.e., greater overall sensitivity to musical reward). This may be because the total BMRQ score reflects various aspects of reward sensitivity, rather than simple pleasure. The BMRQ has five subscales: Music Seeking, Emotion Evocation, Mood Regulation, Sensory-Motor, and Social Reward. Among these, ‘pleasure’ seems to align most closely with Emotion Evocation, which includes questions such as, “I sometimes feel chills when I hear a melody that I like.” However, other subscales, such as Social Reward—which includes items including, “At a concert I feel connected to the performers and the audience”—do not directly measure the pleasure experienced while listening to music. Therefore, the discrepancies between our findings and those of previous studies (Mas-Herrero et al. 2012; Honda et al. 2023) may be attributable to the fact that we specifically measured experienced ‘pleasure’ during music listening, while the total BMRQ score captures a broader concept of musical reward. In addition, while our task assessed state-like aspects of the groove experience, the BMRQ evaluates trait-like characteristics. This distinction may have also contributed to the differences observed in the results. In the older group, we found no sex difference in experienced pleasure. This may be explained by differences in the lifetime change in reward sensitivity between males and females. A study reported that males tend to show a plateau or moderately increasing sensitivity between ages 18 and 29, followed by a decrease from age 30 onward, whereas females generally exhibit a steady decrease in sensitivity beginning before age 18 (Cardoso Melo et al. 2023). As a result, the difference in sensitivity between males and females is most pronounced in younger adults, especially between ages 18 and 29, but diminishes in older age groups. This may explain why we found no significant sex difference in experienced pleasure in the older group. Contrary to the “pleasure” ratings, there was no significant difference in the “wanting to move” ratings between males and females. A study found that girls participated more in extracurricular dance than boys (Anderson et al. 2017), suggesting that females are generally more motivated to engage in dance activities. However, a study investigating dance movement in a club-like environment found that, although females tended to move more intensely, there was no significant difference between males and females (Van Dyck et al. 2013). Therefore, the findings from these studies and our study suggest that females may be more inclined to participate in dance, while there may be no significant difference in intensity or motivation to move between males and females once exposed to music.

There are several limitations to our study. First, it is important to note that NM-MRI CNR reflects long-term dopamine function rather than short-term dopamine function in individuals. NM is produced during dopamine synthesis and gradually accumulates within neurons over time (Zecca et al., 2002), reflecting long-term, trait-like dopamine function. In contrast, PET assesses dopamine synthesis capacity by measuring the uptake of DOPA and its decarboxylation into dopamine by aromatic L-amino acid decarboxylase in dopaminergic neurons (Kumakura & Cumming, 2009), thereby reflecting short-term, state-like dopamine function. Therefore, to evaluate dynamic dopamine function in individuals, PET imaging of the dopamine system is required. This may help overcome the limitation of our study in investigating the relationship between the groove experience and dopamine function in older populations. Second, the small sample size is a notable limitation and may have influenced the results since we were only powered to detect large effects. Future studies should include larger samples to enable more robust conclusions. Finally, it is important to note that, because we examined four different variables (“pleasure” and “wanting to move” ratings in both younger and older groups), we did not apply corrections for multiple comparisons, despite performing four linear mixed-effects regression models. Further research is warranted to draw a robust conclusion regarding the relationship between the groove experience and dopamine function.

## 5. Conclusion

In this study, we investigated the relationship between NM levels measured with MRI (NM-MRI CNR) in the SN and the groove experience. Our results revealed that NM-MRI CNR in the SN was positively related to the “pleasure” aspect of the groove experience in the younger group, while no relationship was found between NM-MRI CNR and the “wanting to move” aspect of the groove experience in either the younger or older groups. These results suggest that the “pleasure” aspect of the groove experience may be related to dopaminergic function in the SN in younger populations, while “wanting to move” maybe not. Our results also indicate that, while previous studies have shown a relationship between the “pleasure” and “wanting to move” aspects (e.g., Düvel et al. 2021; Etani et al. 2018; Senn et al. 2020; Witek et al. 2014), they are not identical and likely involve different dopaminergic pathways, which warrants further studies.

## Acknowledgement

This study was supported by JST PRESTO Grant (JPMJPR23S90), and JST COI-NEXT Grant (JPMJPF2203) awarded to S.F, and by the Japan Society for the Promotion of Science (18H02755, 22H03002), Japan Agency for Medical Research and Development (AMED: JP24wm0625302), Japan Research Foundation for Clinical Pharmacology, Naito Foundation, Watanabe Foundation, and Takeda Science Foundation awarded to S.N. We thank Mr. Nishikata for his technical support.

